# Combined single-sample metabolomics and RNAseq reveals a hepatic pyrimidine metabolic response to acute viral infection

**DOI:** 10.1101/2022.08.26.505340

**Authors:** Zachary B Madaj, Michael S. Dahabieh, Vijayvardhan Kamalumpundi, Brejnev Muhire, Dean J. Pettinga, Rebecca A. Siwicki, Abigail E. Ellis, Christine Isaguirre, Martha L. Escobar Galvis, Lisa DeCamp, Russell G. Jones, Scott A. Givan, Marie Adams, Ryan D. Sheldon

## Abstract

**Objective:** Metabolomics and RNA sequencing (RNAseq) each provide powerful readouts of phenotype, and integration of these data can provide information greater than the sum of their parts. The ability to conduct such analysis on a single sample has many practical advantages, especially when dealing with rare or difficult-to-obtain samples. While methods exist to isolate multiple biomolecular subclasses from the same sample, in-depth analysis of the suitability of these approaches for multi-‘omics readouts is lacking.

**Methods:** Mice were injected with lymphocytic choriomeningitis virus (LCMV) or vehicle (Veh) control and liver tissue was harvested 2.5-days later. RNA was isolated from aliquots of pulverized liver tissue either following metabolite extraction using 80% methanol (MetRNA) or directly from frozen tissue (RNA). RNA sequencing data was evaluated by differential expression analysis via edgeR and dispersion using Gini’s mean differences. Differential metabolite abundance was assessed using LIMMA. Pathway enrichment analysis was conducted on metabolomics and RNAseq data using MetaboAnalyst’s joint-integration tools.

**Results:** Prior metabolite extraction had no deleterious effects on quality or quantity of isolated RNA. RNA and MetRNA generated from the same sample clustered together by principal component analysis, indicating that inter-individual differences were the largest source of variance. Of the 2,169 genes that were differentially expressed between LCMV and Veh, the vast majority (n=1,848) were shared between extraction method, with the remainder evenly divided between RNA (n=165) and MetRNA (n=156). These differentially expressed genes unique to extraction method were attributed to randomness around the false discovery rate (FDR) = 0.05 cutoff and stochastic changes in variance estimation. Gini analysis further revealed that extraction method had no effect on the dispersion of detected transcripts across the entire dataset. To demonstrate the power of multi-omics integration on interrogated metabolic phenotypes, we next performed integrated pathway enrichment analysis on RNAseq data and metabolomics data. Our analysis revealed pyrimidine metabolism as the most impacted pathway by LCMV infection. Plotting up- and down-regulated genes and metabolites on the Kyoto Encyclopedia of Genes and Genomes (KEGG) pyrimidine pathway exposed a pattern enzymatic degradation of pyrimidine nucleotides to generate the nucleobase uracil. Further, uracil was among the most differentially abundant metabolite in serum of LCMV infected mice, suggesting a novel mechanism of hepatic uracil export in acute infection response.

**Conclusions:** We demonstrate that prior metabolite extraction does not have a deleterious effect on RNAseq quality, which enables investigators to confidently perform metabolomics and RNAseq on the same sample. Implementation of this approach revealed a novel involvement of the hepatic pyrimidine metabolism during acute viral infection.

## 1. Introduction

Omics approaches are powerful tools to understand complex biological phenotypes. However, relying on a single ‘omics readout gives an incomplete and at times misleading view of the biological system. This is particularly true when interrogating metabolic phenotypes. Metabolomics, or the measurement of small-molecule metabolite abundances, provides a snapshot in time of the current metabolic state of the system. While metabolomic profiling is useful in identifying metabolites or metabolic pathways involved in a phenotype, it fails to capture metabolic dynamics or directionality and can be difficult to interpret in isolation [1]. Similarly, transcriptomic approaches, such as RNA sequencing (RNAseq), provides a semi-quantitative view of transcript abundance that reflects cellular goals. However, in the context of metabolism, the expression of a gene or its translation to protein does not necessarily equate to the activity of a given metabolic pathway (PMID: 35981545). Therefore, integration of metabolomics and RNAseq datasets can help paint a more complete picture of a particular metabolic phenotype.

Both RNAseq and metabolomics rely on the destruction of the sample from which the target molecules are isolated. The sample must be regenerated or sampled multiple times for additional analyses, introducing sampling error and limiting feasibility in low-quantity or rare sample types. For metabolomics, the wide range of compound chemistries present in the metabolome poses a particular challenge as no extraction method is suitable for the entire metabolome. A widely used approach to address this problem is the homogenization of the sample with ice-cold 80% methanol, which both quenches metabolism through protein precipitation and extracts polar and semi polar metabolites involved in central carbon metabolism [2, 3]. After extraction, the insoluble fraction contains protein, neutral lipids, and nucleic acids. Numerous studies have demonstrated that isolation of these molecules for further analyses is possible with additional processing steps [4-10]. However, these studies do not compare the quality of these other target compounds downstream of metabolite extraction *versus* a traditionally extracted control. As such, it is unknown whether this additional processing influences the downstream ‘omics readout and ultimately the biological interpretation.

To address this gap, the main objective of this study was to determine the effects of prior metabolite extraction on RNAseq. To study the potential effects, we used liver tissue from mice infected with a model of acute lymphocytic choriomeningitis virus (LCMV) infection. The liver is both a master regulator of systemic metabolic homeostasis and an immunological hub owing to its large population of non-parenchymal cells including phagocytes (Kupffer cells, macrophages, dendritic cells, neutrophils), and lymphocytes (innate lymphoid cells, T cell and B cells; reviewed in [11]). LCMV is commonly used to study adaptive immunity as it induces strong systemic effects in many organs, including spleen, brain, kidney, and liver in mice [12]. At the peak of LCMV infection, although the liver bears the second largest viral burden (next to the spleen) it possesses the capacity to clear the infection, while robust viral titers are still observed in brain and kidneys [12].

Thus, the LCMV-infected liver (*versus* non-infected controls) is an excellent model system that features strong metabolic and transcriptional responses, providing rich biological contrast, and that is large enough to be sampled multiple times to allow for direct comparison of extraction methods. Our results demonstrate that prior metabolite extraction does not impact RNAseq quality. We further use integrated single-sample metabolomics and transcriptomics data that supports a novel role for pyrimidine metabolism in the hepatic response to LCMV infection.

## 2. Material and Methods

### 2.1. Animal Care/Mice

Twenty male-C57/BL6 mice were selected for the study. All mice were maintained in a climate-controlled environment on a 12h:12h light-dark cycle with *ad libitum* access to water and standard chow diet. At eleven weeks of age, mice were randomly selected to receive an intraperitoneal injection of lymphocytic choriomeningitis virus (LCMV) (Armstrong strain, 2×10^5^ PFU by intravenous injection) or PBS vehicle (Veh) [13]. 2.5 days post-infection, mice were anesthetized with an isoflurane vaporizer. While anesthetized, the liver was removed and snap-frozen in liquid N_2_ for later processing. All animal protocols were approved by the Van Andel Institute Institutional Animal Care and Use Committee.

### 2.2. Liver processing

Frozen livers were pulverized to a fine powder using a mortar and pestle pre-chilled in liquid N_2_. Being careful to avoid thawing of powdered liver, two roughly 30 mg aliquots of each liver were rapidly weighed into liquid N_2_ chilled tubes. One aliquot was weighed into a liquid N_2_ chilled 1.5 mL Eppendorf tube for later RNA extraction. The other aliquot was weighed in to a 2.0 mL Omni Bead Ruptor (19-627, Omni) tube for metabolite extraction. Precise weights were then recorded, and samples were stored at –80°C until processing.

### 2.3. Metabolite extraction

Metabolites were extracted from powdered liver aliquots by homogenization in 80% methanol (v/v) using a bead mill homogenizer in a liquid N_2_ chilled homogenization chamber to prevent sample warming. Extraction solvent volume was adjusted for each sample to achieve a fixed 40 mg tissue/mL extraction solvent ratio. After homogenization, samples were incubated on wet ice for 60 minutes to ensure complete metabolite extraction and precipitation of protein and nucleic acids. Extracts were then centrifuged at 17,000 *x g* for 10 minutes at 4°C. The supernatant was transferred to a fresh 1.5 mL Eppendorf tube, dried in a Speedvac (Genevac) and stored at -80°C for later LC/MS analysis. The insoluble pellet was also dried and stored at -80°C for RNA extraction.

### 2.4. Metabolomics

Metabolite profiling analysis was completed using ion-paired reversed phase liquid chromatography using an HPLC (1290 Infinity II, Agilent Technologies, Santa Clara, CA) coupled to a triple quadrupole mass spectrometer (6470, Agilent) with electrospray ionization operated in negative mode. The column was a ZORBAX Rapid Resolution HD (2.1 × 150 mm, 1.8 µm pore size; 759700-902, Agilent). Mobile phase A was 3% methanol (in H_2_O), and mobile phase B was 100% methanol. Both mobile phases contained 10 mM of the ion-pairing agent tributylamine (90780, SigmaAldrich, St. Louis, MO, USA), 15 mM acetic acid, and 2.5 µM medronic acid (5191-4506, Agilent Technologies, Santa Clara, CA, USA). The LC gradient was: 0-2.5min 100% A, 2.5-7.5 min ramp to 80% A, 7.5-13 min ramp to 55% A, 13-20 min ramp to 99% B, 20-24 min hold at 99% B. Flow rate was 0.25 mL/min, and the column compartment was heated to 35°C. The column was then backflushed with 100% acetonitrile for 4 minutes (ramp from 0.25 to 0.8 mL/min in 1.5 minutes) and re-equilibrated with mobile phase A for 5 minutes at 0.4 mL/min. Differential abundance of metabolites collected from liver tissue and blood serum were analyzed separately using the same R v 4.1.0 (https://cran.r-project.org/) workflow. To start, metabolites with zero variance were filtered out, then the remaining metabolites were assessed for missingness. Metabolites with missing data were determined to be not-missing-at-random as these metabolites had lower overall mean abundance compared to metabolites without any missingness. To address this potential source of bias, metabolites with less-than 30% missingness were imputed from a left truncated distribution with parameters estimated using quantile regression (https://cran.rstudio.com/web/packages/imputeLCMD/index.html). Metabolite data were then log2 transformed and a variance stabilizing normalization was applied to prepare data for analysis via LIMMA ebayes; both the limma linear regression and empirical bayes were fit using robust methods to prevent individual animals and/or metabolites from exerting excessive influence on estimates (https://bioconductor.org/packages/release/bioc/html/limma.html). Batch was included as covariate in the LIMMA linear models. Hypothesis tests were then multiple-testing adjusted via Benjamini-Hochberg to maintain a 5% false discovery rate (FDR).

### 2.5. RNA Extraction and Quality Control

RNA was isolated directly from frozen liver tissue (RNA) or from post-metabolite extracted liver tissue (MetRNA). Samples were pulverized using QIAzol Lysis Reagent and a Tissue Lyser II. RNA was then purified from the lysates using RNeasy Plus Universal Mini Kit column-based isolation per manufacturers’ instructions. Isolated RNA was quantified by Promega Quantus and measured for integrity using Agilent Bioanalyzer. We used the calculated RNA Integrity number (RIN), which used the entire electrophoretic trace of RNA sample (including degradation products) to compare the RNA integrity between our samples. Differences in unique mapping rates between LCMV vs Veh and MetRNA vs RNA were tested using beta regression via the R package ‘betareg’ (https://cran.r-project.org/web/packages/betareg/index.html).

### 2.6. RNA Sequencing

Following quality control, stranded RNA libraries were prepared from 500 ng RNA using the KAPA - mRNA Hyper Prep (08098140702, Roche). RNA libraries were then normalized, pooled, and sequenced on Illumina NovaSeq S1 flow cell using paired end, 50 base pair reads to an average depth of 50 M reads per sample. Base calling was performed by Illumina RTA3 and output of NCS was demultiplexed and converted to FastQ format with Illumina Bcl2fastq v1.9.0.

Adapters and low-quality bases (Phred>30) were trimmed from raw reads using Trim_Galore (https://www.bioinformatics.babraham.ac.uk/projects/trim_galore/). These trimmed reads were then aligned to the GRCm38 reference genome using the STAR aligner v2.7 (https://pubmed.ncbi.nlm.nih.gov/23104886/). Aligned read counts were imported into edgeR (https://doi.org/10.1093/bioinformatics/btp616) wherein library sizes were normalized using the trimmed mean of M values method and pairwise comparisons for differential expression were performed using the quasi-likelihood approach as in (https://f1000research.com/articles/5-1438/v2).

### 2.7. Statistical Analysis of RNA Dispersion

To determine if isolating RNA and metabolites from the same sample altered variance profiles within the RNAseq gene expression estimates (e.g., does coextraction introduce additional noise into the measurements), we compared Gini’s mean differences (Gini’s MD) between samples where only RNA was isolated and samples where RNA was isolated post-metabolite extraction, stratifying them by experimental group (LCMV/Veh) (https://cran.r-project.org/web/packages/Hmisc/index.html). Gini’s MD was chosen as the measure of dispersion because it is non-parametric and has a natural interpretation. The distributions of the fold-change in dispersion were then assessed for skewness which could indicate a biased change in dispersion related to the extraction method, particularly if this bias is conserved over both LCMV and vehicle groups. Finally, Gini’s MD from the two extraction methods were correlated using Pearson’s rho.

### 2.8. Pathway Enrichment Analysis & Multi-omics

Multi-omic pathway enrichments were performed using MetaboAnalyst’s v5.0-‘joint-pathway analysis’ where we investigated which metabolic pathways were most impacted by LCMV infection (https://www.metaboanalyst.ca/). All differentially expressed genes and differentially abundant metabolites with less-than a 5% FDR were included in the enrichment analysis; gene expression fold-change was also considered. Pathway impact and Benjamini-Hochberg FDR adjusted significance were estimated using hypergeometric tests and MetaboAnalyst’s ‘tight integration’ method, which combines queries into one pooled universe. The betweenness centrality was emphasized to identify pathways where the flow of information was heavily impacted. The ‘overall weighting’ and‘pathway-level’ weighting methods were also explored in combination with each centrality measures (degree, betweenness, and closeness). MetaboAnalyst was also used to examine individual impacts of DEGs and DAMs on all pathways and metabolic pathways, respectively. The top 5 impacted pathways are reported. A more common gene set enrichment analysis via the R package clusterProfilier [14] and reactome pathways was also conducted for comparison.

## 3. Results

### 3.1. RNA quality

First, we assessed whether there were any overt differences in RNA quality from prior metabolite extraction. Both methods yielded RIN scores >9 and a 260/280 ratio of ≈2 (Figure S1A-B). MetRNA had an increased 260/230 ratio (Figure S1C), which could be an indication of removal of contaminants, and improved RNA yield (Figure S1D). Thus, the metabolite extraction carried out prior to the RNA purification does not compromise RNA quality and might serve to improve contaminate removal and RNA yield.

### 3.2. RNAseq analysis

RNAseq was performed on both RNA and MetRNA samples. Unique mapping rates averaged 75% for vehicle and 82% for LCMV treated samples; samples were diverse and expressed 15,535 unique gene features after trimming. Principal component analysis (PCA) attributed 65% of variation in the dataset to treatment (LCMV *vs* Veh, p=0.0001). RNAseq data from each extraction approach were compared. Interestingly, the unique mapping rate, i.e., the number of sequenced fragments that map to a unique location in the genome, was elevated in LCMV *vs* Veh (p < 0.0001), and in MetRNA *vs* RNA (p = 0.02) (data not shown). Elevation in LCMV over vehicle was hypothesized to be due to gene activation in response to viral infection, while the unique mapping rate of MetRNA correlates to the improved 260/230 ratio and supports the indication that the metabolite extraction may have improved contaminant removal.

We next asked whether prior metabolite extraction affected the detection of significantly differentially expressed genes between biological groups. Because of the increased processing steps in MetRNA, we hypothesized the intra-group variance of these samples may increase, and consequently the number of expressed genes that are significantly different between the two groups would decrease. We found that the majority of differentially expressed genes (1,848) between LCMV and Veh were shared between extraction methods (**Figure 1G**). There were 321 differentially expressed genes that were significant in only one extraction method, with 165 unique to RNA and 156 unique to MetRNA (**Figure 1G**). A delta-delta comparison of these genes [LCMV_MetRNA – Veh_MetRNA] - [LCMV_RNA – Veh_RNA] produced no significant findings (all FDR > 0.999). The median fold-change between genes uniquely significant to an extraction method was 1.04 with 94% of the genes having a log2 (fold change) less than +/- 1. The probability of obtaining this amount of overlap (>85% shared DEGs) by chance, as estimated via a hypergeometric test, is < 0.0001. Further, as seen in **Figure 1H**, the two methods are highly concordant with a Pearson’s R^2^ = 0.97. Taken together these analyses suggest that alterations in the detection of significantly differentially expressed genes by extraction method is due to stochastic differences in measurements, rather than a systematic effect of extraction.

**Figure 1:**
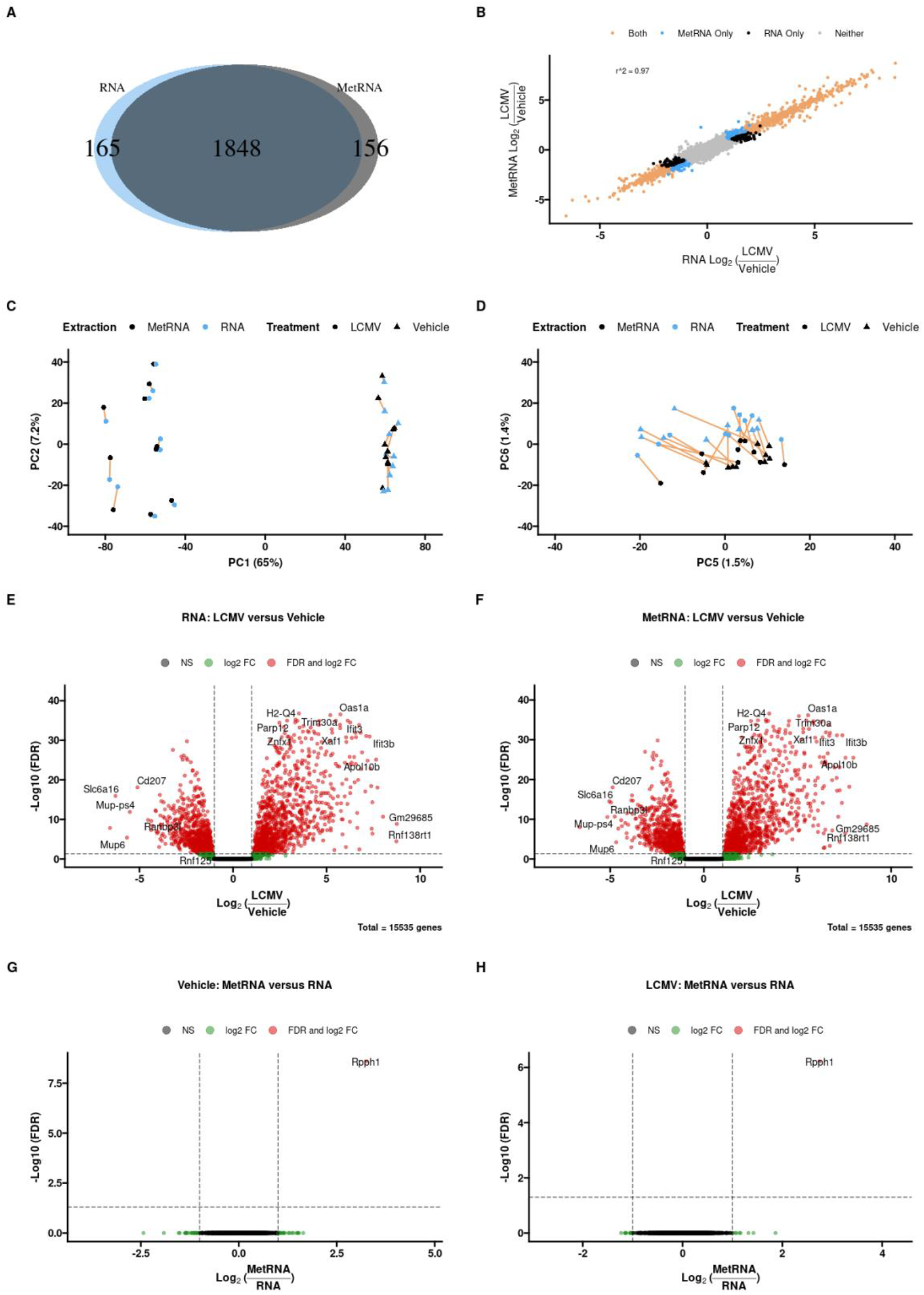
Comparison of the RNAseq data obtained with the two extraction methods (RNA: classical mRNA isolation; MetRNA: metabolite extraction was performed prior to mRNA isolation). A) Venn diagram of the genes found to be differentially expressed with FDR < 0.05 in both extraction methods, <85% of the 2,169 genes found to be differentially expressed in at least one extraction method were significantly different in both. B) Correlation of the log2 fold-change in gene expression (LCMV/Veh) between the two extraction methods. Over 97% of the variance in one extraction method was explained by the other, further genes that were only identified as differentially expressed in one method had only slight changes in estimated fold-change. C) The first two principal components show distinct LCMV and vehicle clusters. Data do not cluster by extraction method, but rather by sample ID (LCMV/Veh). D) The fifth and sixth principal components show separation by extraction method. PC6 is comprised predominantly by RPPH1. E) and F) volcano plots of genes differentially expressed between vehicle and LCMV animals within both extraction methods. G) and H) volcano plots of genes differentially expressed between the extraction methods within vehicle and LCMV animals, respectively.

Scores plots from principal component analysis (PCA; **Figure 1A-B**) revealed robust separation between biological groups in principal component 1 (PC1, 65% of variance), followed by intragroup differences in PC2 (7.2% of variance; **Figure 1A**). Separation on PCA score plot by extraction method was only apparent in PC5 and PC6 (1.5% and 1.4% of variance, respectively; **Figure 1B**). Volcano plots comparing LCMV/Veh gene expression in RNA (**Figure 1C**) and MetRNA **(Figure 1D)** extraction methods revealed nearly identical distributions. Only a single gene, *Rpph1*, was significantly differentially expressed by extraction method in both Veh (**Figure 1E**) and LCMV (**Figure 1F**) groups. *Rpph1* is a highly conserved lncRNA that encodes the RNA component of RNaseP, which processes tRNA precursor transcripts. In eukaryotes, RNaseP exists as a riboenzyme with up to ten protein subunits [15]. In light of our observations that MetRNA improves total RNA yield (Figure S1), the fact that we observed an increase in Rpph1 expression might indicate that the additional steps prior to RNA extraction improved the extraction of protein-complexed RNAs.

### 3.3. Extraction method does not affect data dispersion

Having demonstrated that prior metabolite extraction does not affect the distribution of differentially expressed genes, we next sought to understand what effects extraction method has on global dispersion of RNA. That is, does coextracting metabolites and RNA introduce or reduce noise? To explore this question, we utilized Gini’s mean difference, a non-parametric estimation of dispersion. Within both LCMV and vehicle-injected animals, Gini’s MD was significantly correlated between MetRNA and RNA only transcripts (both *p*-values were < 0.001; **Figure 2A, 2B**). Over 85% of the variance in the gene expression of vehicle injected animals was conserved between extraction methods, and over 90% in LCMV injected animals. We then examined the distribution of log2 fold-change in Gini’s MD for both LCMV and vehicle animals (**Figure 2C, 2D**); both distributions appeared Gaussian with skewness estimates of only ∼0.1, LCMV mean = 0.04, vehicle mean = –0.04, LCMV median= 0.05 and vehicle median= -0.04. These low skewness estimates, combined with means approximately equal to medians strongly suggest that the changes in dispersion are symmetric about a null difference (i.e., 1-fold difference or log2(fold) = 0), and therefore random.

**Figure 2.**
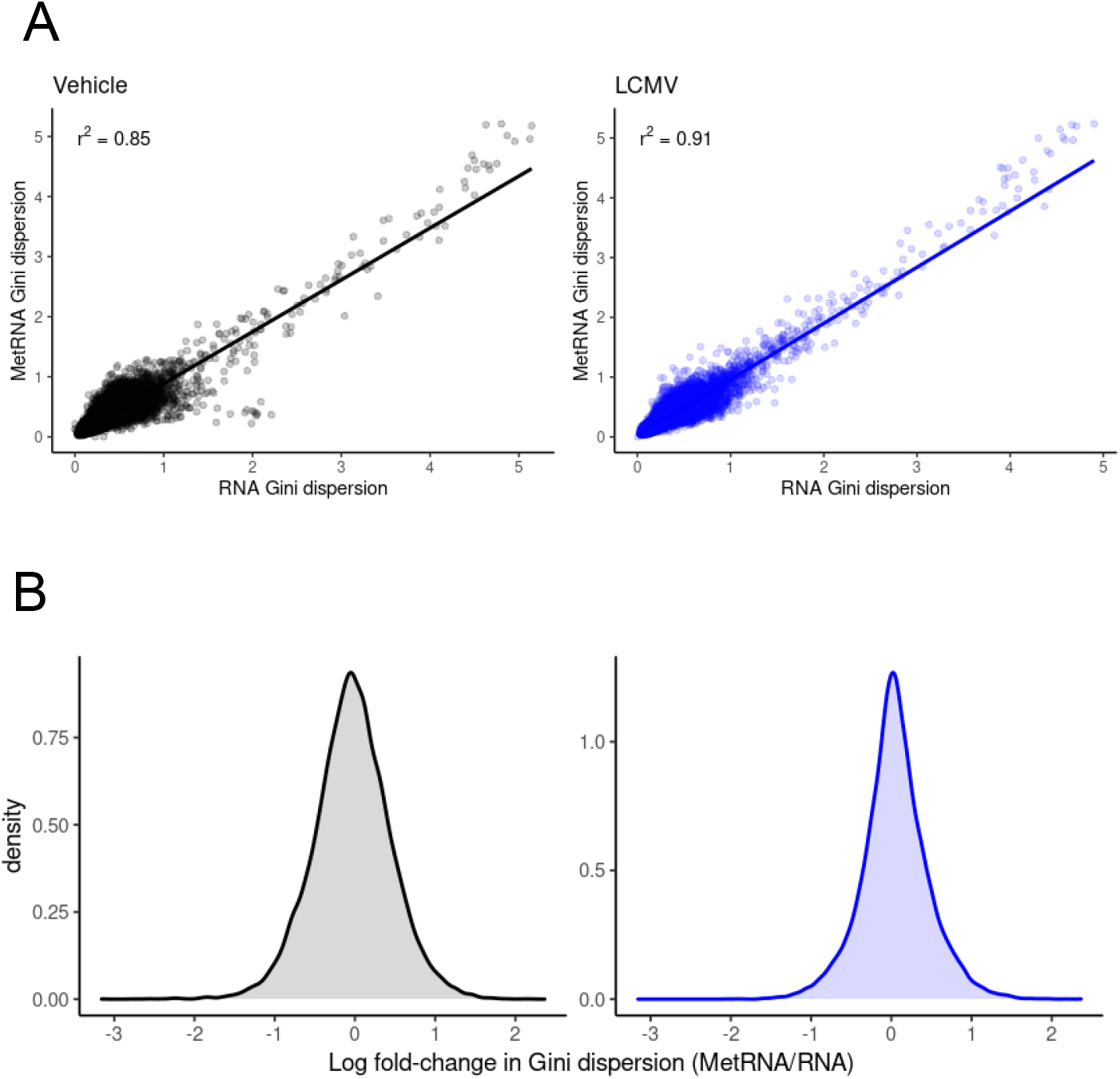
Analysis of Dispersion A) Correlating Gini’s MD of each gene between the two extraction methods in vehicle and LCMV animals; both were significantly correlated (p < 0.05). B) Estimated densities of the fold-change in Gini’s MD between the extraction methods in LCMV and vehicle animals; both are roughly symmetric about 0.

### 3.4. Integrated pathway analysis

Our data support the conclusion that RNAseq from post-metabolite extraction RNA was at least as informative as traditionally extracted RNA, which should enable confident multi-omics approaches to understanding metabolic phenotype. To evaluate the power of this approach, we interrogated metabolic phenotypes using combined RNAseq and metabolomics from the same sample. As with the broad effects of LCMV on gene expression (**Figure 1**), we observed that LCMV induced broad changes in the hepatic metabolome (Figure S2). Pathway enrichment analysis of metabolomics data revealed purine, pyrimidine, and glycolysis/gluconeogenesis as the most differentially impacted pathways (Figure S3). However, within a given pathway some metabolites had increased, and others decreased. This leaves ambiguity as to how these pathways are affected with LCMV.

To gain further insight, we next performed pathway enrichment analysis on RNAseq data. Unsurprisingly, the results of this analysis, as generated by both MetaboAnalyst and the more common GSEA method, were dominated by immune response pathways in LCMV infected mice (Figure S4). This revealed JAK-STAT signaling pathway, linoleic acid metabolism, beta-Alanine metabolism, neutrophil degranulation, signaling by interleukins, and cytokine signaling in the immune system as some of the most impacted pathways. Interestingly, there was little overlap between RNAseq metabolic pathways, and those pathways identified with metabolomics data.

Finally, we integrated metabolite and RNA data and performed a combined pathway analysis (**Figure 3)**, revealing pyrimidine metabolism (Kegg pathway map 00240) as the most highly impacted pathway (**Figure 3A**). Pyrimidine metabolism was consistently a significantly and heavily impacted pathway under each of the various centrality measures and weighting methods used for the joint integration, often remaining the most affected (Figure S5). Both transcripts and metabolites in the pyrimidine metabolism pathway had bidirectional responses to LCMV (**Figure 3B, 3C**), which obscured biological interpretation. First, nucleotides (CTP, CMP, UTP, UDP, UMP, dTTP, dTDP) were decreased in LCMV, whereas metabolites involved in *de novo* pyrimidine synthesis (*N*-carbamoyl-aspartate, dihydroorotate, orotate), nucleosides (cytidine, uridine, thymidine, deoxycytidine, deoxyuridine, thymidine), and nucleobases (uracil, thymine) were increased (**Figure 3C**). When considering metabolomics data in isolation, it is clear that pyrimidine metabolism is involved in the phenotype. However, it is unclear whether *de novo* nucleotide production is elevated to support increased RNA/DNA synthesis, or whether nucleotides are being catabolized to produce nucleosides/nucleobases.

**Figure 3.**
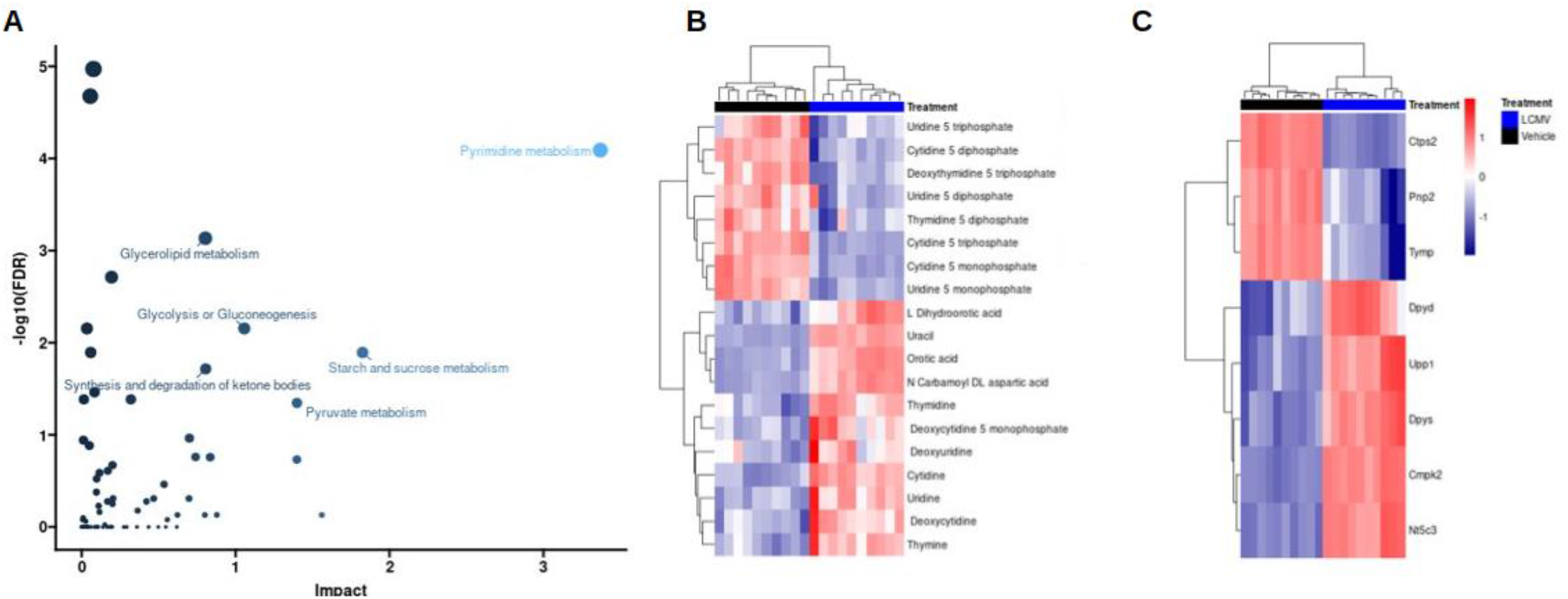
Integrated Metabolic Pathway Enrichment. A) Plot of how much each pathway examined by MetaboAnalyst was impacted by changes in gene expression and whether or not these changes were significant after FDR adjustment. Data are colored by impact and sized by FDR values. The pyrimidine metabolism pathway was the most impacted (FDR < 0.0001). B) and C) heatmaps of pyrimidine metabolites and transcripts that were measured in this study. For both –omics, we observed both up and down regulation within the pathway and strong clustering within LCMV/Vehicle.

We next asked whether gene expression data could inform directionality of the pyrimidine metabolite phenomena. We mapped gene expression and metabolite heatmap signatures onto the KEGG pyrimidine pathway (**Figure 4A**). RNAseq revealed that the expression of genes that encode enzymes catalyzing bidirectional nTP ↔ nDP (2.7.4.6), nDP ↔ nMP (2.7.4.14), and nucleoside ↔ nucleobase (2.4.2.3) interconversions were either unchanged or increased. Also, expression of genes encoding proteins that drive dephosphorylation of nucleotide monophosphates to the cognate nucleoside were increased. In contrast, genes encoding enzymes involved in nucleobase catabolism (1.3.1.2 and 3.5.2.2) had decreased expression. To further highlight this pattern, cytidine nucleotides (CTP, CDP, CMP) have decreased abundance (**Figure 4B**), whereas the gene (*Nt5c3*) encoding the cytosolic 5’-nucleotidase 3 had elevated expression (**Figure 4C**) and the product of this reaction, cytidine, was also elevated (**Figure 4B**). Finally, the pyrimidine nucleobase, uracil, was among the most elevated metabolites in the liver of LCMV infected mice (**Figure 5A**). We also conducted plasma metabolomics in these mice. In line with our observations of increased hepatic uracil content, plasma metabolomics revealed that uracil, along with the inflammatory mediator metabolite itaconate [16, 17], were the most strongly induced metabolites in plasma upon LCMV infection (**Figure 5B**). Together our data suggest that pyrimidine nucleobases, especially uracil, are being generated in the liver through the catabolism of pyrimidine nucleosides from both *de novo* pyrimidine synthesis and pyrimidine nucleotide catabolism and that they are potentially released into the circulation.

**Figure 4.**
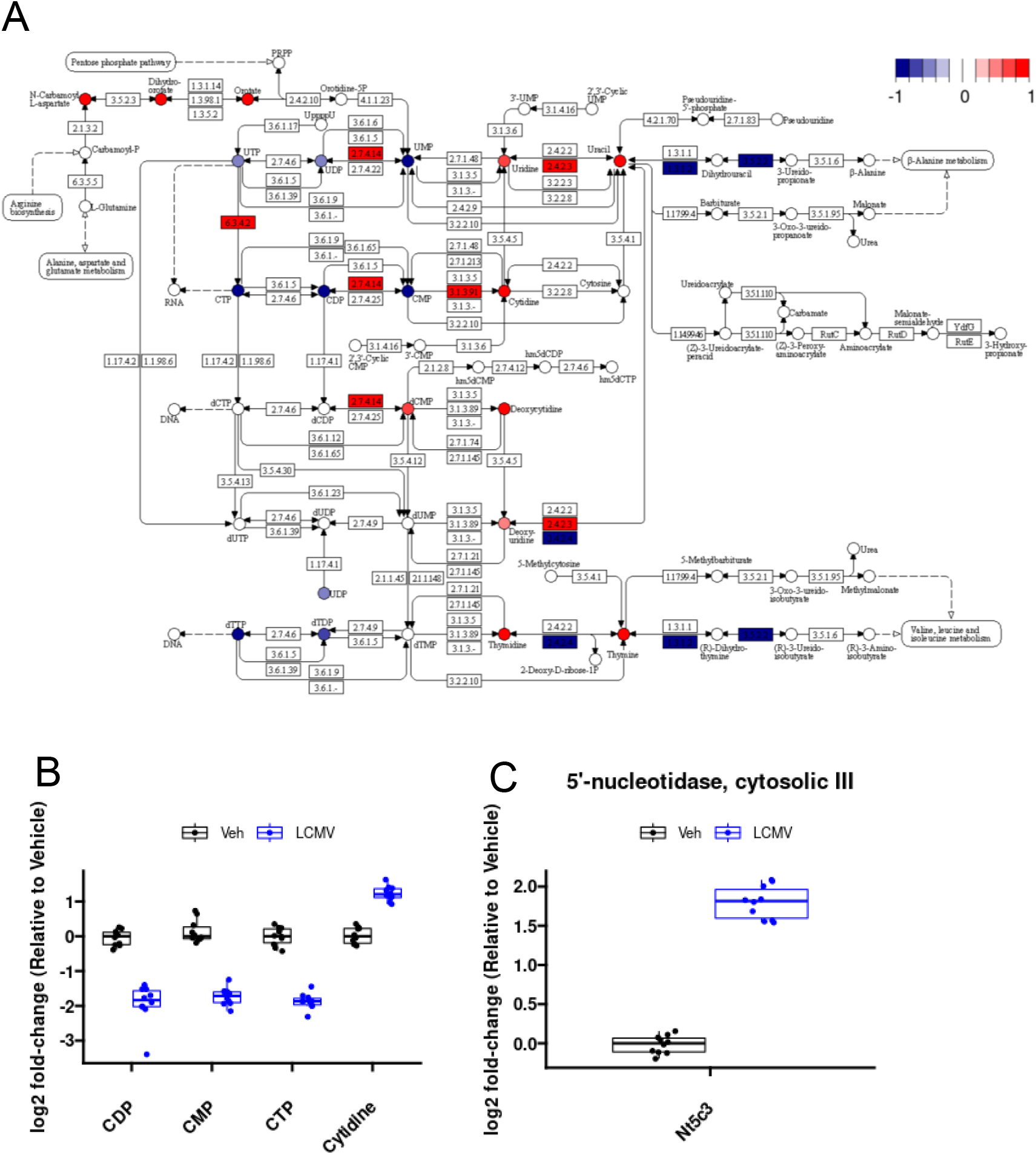
Integrative multi-omics analysis demonstrating biological information gain. (A) Significantly different genes and metabolites colored by log2 fold-change mapped to KEGG pathway. (B) Cytidine metabolites differentially abundant in the liver. (C) Differential expression of 5’-Nucleotidase, Cytosolic IIIA in the liver.

**Figure 5.**
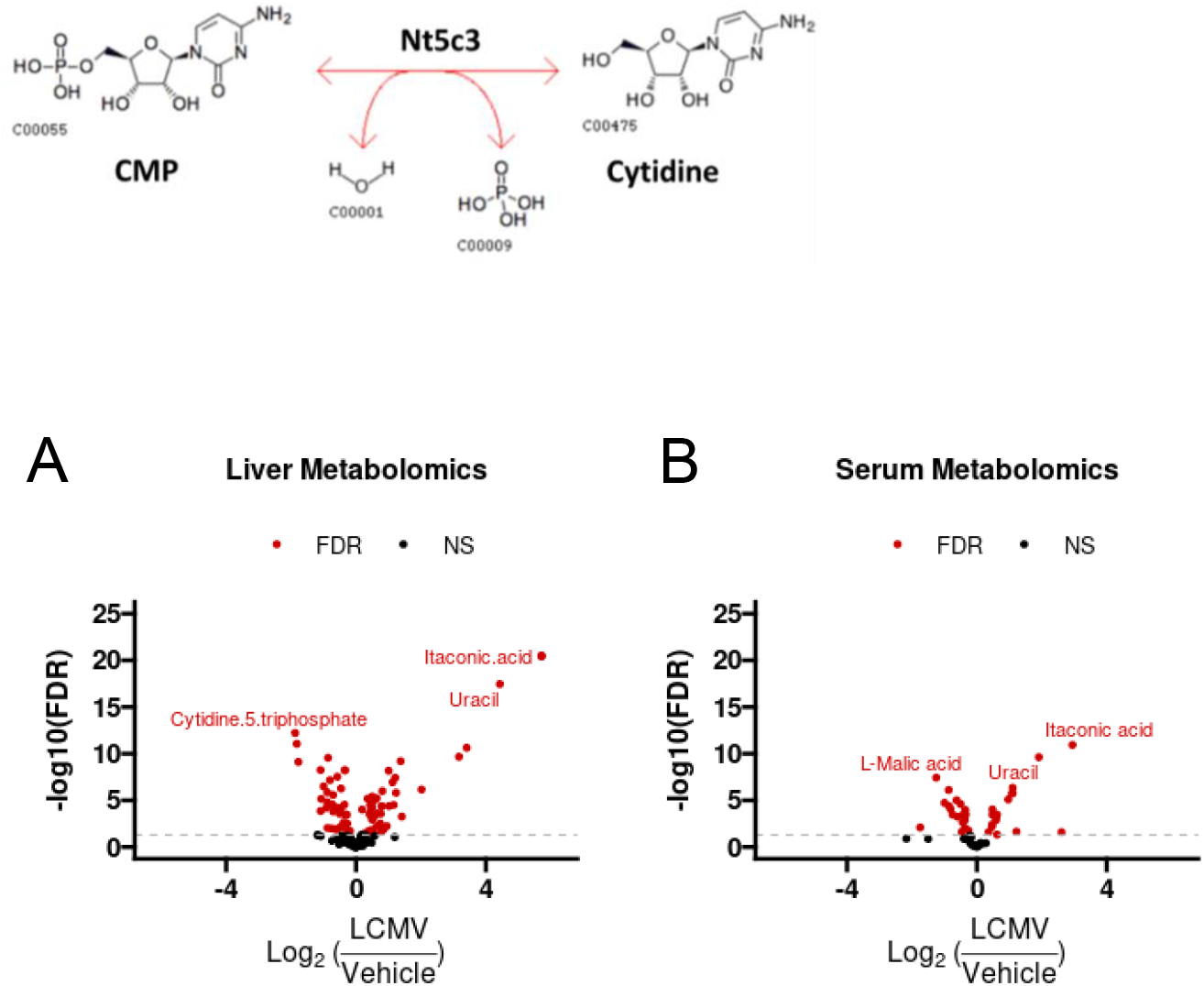
Volcano plots of metabolomics data in the (A) liver and (B) serum reveal strong induction in uracil abundance. “FDR”: Indicates Benjamini-Hochberg false discovery rate adjusted *p*-values < 0.05. “NS”: not significant.

## 4. Discussion

Conducting multiple ‘omics approaches on a single sample provides a more complete picture of the phenotype while avoiding challenges like repeat-sampling and limited quantities but it also requires the serial extraction of different classes of biomolecules. While numerous such protocols are available [4-10], there is a lack of information on whether and how the readout is affected by additional processing steps. In this study, we used liver tissue from vehicle or acute LCMV infected mice to assess the effects of prior metabolite extraction on RNAseq data. Our results demonstrate that RNAseq analysis following metabolite extraction yields virtually identical results as traditionally extracted RNA, not only in data dispersion but most importantly in detection of biological effects. These data provide important and needed validation that co-extraction of metabolites and RNA is a viable approach that is at least as effective as standard approaches that gain a single outcome per sample.

A major advancement of this work is the direct and comprehensive evaluation RNAseq data quality from RNA isolated following metabolite extraction compared to samples only undergoing RNA isolation. We took several steps to allow direct comparison extraction methods rather than sampling variability. First, we used snap-frozen mouse liver tissue and pulverized it to a fine powder in liquid N_2_ to avoid liver regional heterogeneity and allow for repeat sampling with minimal variability. Next, to avoid potential errors induced by extraction efficiency, we fixed the tissue weight to extraction volume ratio for each sample. Finally, we selected the LCMV model system that because of its strong biological phenotype, permitted a comprehensive evaluation of extraction method on a complex biological readout.

RNAseq data is commonly interpreted using pathway enrichment analysis, which evaluates the number of significantly differentially expressed genes in a given pathway that occur above what would be expected by chance. While this approach can lead to important insight into pathways that may be involved in the phenotype, establishing how that pathway is changing can be challenging. This is especially true for inferring changes in metabolic activity from RNAseq data because metabolic genes can change in response to the metabolite milieu but also be a cellular response to alter metabolism. Similarly, metabolite pool sizes may indicate that a certain pathway is involved, but directionality is unclear. For example, increased abundance of a metabolite could either mean its production is increased or its clearance is decreased. This is further complicated by the biased nature of metabolomics. The wide range of metabolite chemistries and abundances make truly unbiased sampling of the metabolome impossible. Therefore, metabolic pathway involvement is skewed to those in which metabolites are detectable by the analytical method.

Here, we demonstrate that combining RNAseq and metabolite data can lead to a more complete picture of the metabolic phenotype and allow for the generation of actionable hypotheses. Our results support the hypothesis that LCMV infection induces a liver metabolic program to produce and secrete uracil into circulation. Specifically, levels of CTP, CDP, and CMP, were found to be decreased in liver extracts upon LCMV infection, while cytidine levels were increased (**Figure 4B**). These observations are supported by increased transcript levels of *Nt5c3*, which encodes for the enzyme that dephosphorylates CMP and UMP to their nucleosides cytidine and uridine, respectively [18]. Furthermore, expression of the enzyme that converts uridine to uracil, uridine phosphorylase (Upp1) [19], is upregulated upon LCMV infection (**Figure 3B**), and uracil levels are elevated in both liver and serum in LCMV extracts (**Figure 5**). Thus, while both RNAseq and metabolomics datasets indicated an involvement of pyrimidine metabolism, their integration clearly indicates that pyrimidine nucleotides are being catabolized to support hepatic uracil release. We speculate that a physiological basis for hepatic uracil production during acute infection may be to supplement activated and proliferating immune cells with nucleotides imperative for RNA synthesis. Further investigation is necessary to test this model, but this example highlights the power of integrating multiple ‘omics data to generate an actionable hypothesis.

In our multi-omic pathway analysis we focused on integrating the metabolome and transcriptome data using MetaboAnalyst’s tight integration of the queries and betweenness centrality. Under this integration method the multi-omics results are pooled into a single query where every hit has the same weight. With 36 times more differentially expressed genes than differentially abundant metabolites, this tight integration can result in transcriptomic differences dominating the results. We found strong metabolite evidence of pyrimidine metabolism being impacted in this study. If there were no transcript evidence, we would not expect pyrimidine metabolism to remain one of the top hits in the joint analysis. However, since we did observe differences in the regulation of genes related to pyrimidine metabolism, the result was conserved. This yielded higher confidence that the pyrimidine pathway was truly impacted. The betweenness centrality identifies compounds/transcripts that act as ‘brokers’ or ‘bridges’ between subunits of a pathway. In the context of pyrimidine metabolism, when compounds/transcripts with high betweenness centrality are altered, it indicates that the entire cascade of biochemical signaling has been fundamentally altered. For completeness, we explored other centrality measures and weighted integration methods, each of which indicated pyrimidine metabolism as a significantly impacted pathway (Figure S5). The concordance of these measures provides confidence that infection with LCMV results in a strong involvement in pyrimidine metabolism.

As pyrimidine metabolism was the most significantly impacted pathway in our multi-omic analysis, we used it as an example to highlight the power of multi-omic integration. However, it is important to note that other metabolic pathways were also significantly affected. Joint-integration methods revealed other candidate pathways to explore, notably “Starch and Sucrose Metabolism” and “Degradation of Ketone Bodies”, which were only identified as being highly impacted in the joint analysis.

A general limitation of metabolomics is incomplete coverage of the metabolome. While RNAseq data can provide a comprehensive view of transcripts above a limit of detection threshold, metabolomics coverage is dependent on both analytical methods (metabolite extraction technique, chromatography, ionization type and polarity, and instrument sensitivity) and biological abundance. Therefore, it is important to consider that metabolomics studies, and subsequent multi-omics integration, will be biased towards the metabolites that are detectable on the method. In this light, metabolomics can be used positively, for example to provide orthogonal validation to RNAseq results or highlight altered metabolic pathways independent of transcription. However, the use of metabolomics data to rule out certain metabolic pathways, is not recommended because metabolic pathways are often incompletely covered or simply not detected. By extension, RNAseq results may be used to direct follow-up metabolomics analyses that target specific pathways/compounds of interest.

## 5. Conclusion

Often biological material is limited and the ability to maximize information gained from the same sample is beneficial. Even when tissue is in abundance, regional heterogeneity within a tissue can limit the utility of multi-omics approaches. Our study demonstrates that sequentially extracting metabolites and RNA for metabolomics and transcriptomic analyses does not fundamentally impact gene transcript abundance. We also show that by jointly integrating changes to both –omics’ datasets through enrichment analyses, we can identify metabolic phenotypes with higher statistical confidence, while gaining additional biological insight into how pathways are altered. Finally, through the implementation of these methods we reveal the novel finding that infecting mice with LCMV results in significant and dramatic changes to pyrimidine metabolism. This work provides validation that sequential extraction of metabolites and RNA from a single sample can be confidently used for multi-omics analysis.

## Funding

RGJ is supported by the Paul G. Allen Frontiers Group Distinguished Investigator Program, the National Institute of Allergy and Infectious Diseases (NIAID, R01AI165722), and Van Andel Institute (VAI).

## Competing Interests

RGJ is a scientific advisor for Agios Pharmaceuticals and Servier Pharmaceuticals and is a member of the Scientific Advisory Board of Immunomet Therapeutics.

## Figure and Table Legends

**Table 1:** Pathway enrichment based on the joint-integration of the transcript and metabolite differences. Only the top 5 hits are shown. The ‘Total’ column represents the number of compounds and transcripts queried for a given pathway. ‘Expected’: number of hits one would expect in a given pathway by chance. ‘Hits’: the number of differentially expressed genes and differentially abundant metabolites in each pathway. “FDR”: Benjamini-Hochberg false discovery rate adjusted *p*-values for enrichment based on hyper-geometric tests, and tight integration of the queries using the betweenness centrality. ‘Impact’: the estimated cumulative impact on the pathway.

|  | Total | Expected | Hits | FDR | Impact |
| --- | --- | --- | --- | --- | --- |
| Pyrimidine metabolism | 99 | 11 | 27 | <0.0001 | 3.4 |
| Starch and sucrose metabolism | 37 | 4.1 | 11 | 0.01 | 1.8 |
| Pyruvate metabolism | 45 | 5.0 | 11 | 0.045 | 1.4 |
| Glycolysis or Gluconeogenesis | 61 | 6.8 | 16 | 0.007 | 1.1 |
| Synthesis and degradation of ketone bodies | 10 | 1.1 | 5 | 0.02 | 0.81 |

## References

[1] Jang, C., Chen, L., Rabinowitz, J. D., 2018. Metabolomics and Isotope Tracing, Cell. 173:822–837.

[2] Maharjan, R. P., Ferenci, T., 2003. Global metabolite analysis: the influence of extraction methodology on metabolome profiles of Escherichia coli, Anal Biochem. 313:145–54.

[3] Dettmer, K., Nurnberger, N., Kaspar, H., Gruber, M. A., Almstetter, M. F., Oefner, P. J., 2011. Metabolite extraction from adherently growing mammalian cells for metabolomics studies: optimization of harvesting and extraction protocols, Anal Bioanal Chem. 399:1127–39.

[4] Leuthold, P., Schwab, M., Hofmann, U., Winter, S., Rausch, S., Pollak, M. N., et al., 2018. Simultaneous Extraction of RNA and Metabolites from Single Kidney Tissue Specimens for Combined Transcriptomic and Metabolomic Profiling, J Proteome Res. 17:3039–3049.

[5] Valledor, L., Escandon, M., Meijon, M., Nukarinen, E., Canal, M. J., Weckwerth, W., 2014. A universal protocol for the combined isolation of metabolites, DNA, long RNAs, small RNAs, and proteins from plants and microorganisms, Plant J. 79:173–80.

[6] Roume, H., Heintz-Buschart, A., Muller, E. E., Wilmes, P., 2013. Sequential isolation of metabolites, RNA, DNA, and proteins from the same unique sample, Methods Enzymol. 531:219–36.

[7] Kang, J., David, L., Li, Y., Cang, J., Chen, S., 2021. Three-in-One Simultaneous Extraction of Proteins, Metabolites and Lipids for Multi-Omics, Front Genet. 12:635971.

[8] Salem, M., Bernach, M., Bajdzienko, K., Giavalisco, P., 2017. A Simple Fractionated Extraction Method for the Comprehensive Analysis of Metabolites, Lipids, and Proteins from a Single Sample, J Vis Exp.

[9] Nicora, C. D., Sims, A. C., Bloodsworth, K. J., Kim, Y. M., Moore, R. J., Kyle, J. E., et al., 2020. Metabolite, Protein, and Lipid Extraction (MPLEx): A Method that Simultaneously Inactivates Middle East Respiratory Syndrome Coronavirus and Allows Analysis of Multiple Host Cell Components Following Infection, Methods Mol Biol. 2099:173–194.

[10] Sapcariu, S. C., Kanashova, T., Weindl, D., Ghelfi, J., Dittmar, G., Hiller, K., 2014. Simultaneous extraction of proteins and metabolites from cells in culture, MethodsX. 1:74–80.

[11] Cheng, M. L., Nakib, D., Perciani, C. T., MacParland, S. A., 2021. The immune niche of the liver, Clin Sci (Lond). 135:2445–2466.

[12] Wherry, E. J., Blattman, J. N., Murali-Krishna, K., van der Most, R., Ahmed, R., 2003. Viral persistence alters CD8 T-cell immunodominance and tissue distribution and results in distinct stages of functional impairment, J Virol. 77:4911–27.

[13] Badovinac, V. P., Porter, B. B., Harty, J. T., 2002. Programmed contraction of CD8(+) T cells after infection, Nat Immunol. 3:619–26.

[14] Wu, T., Hu, E., Xu, S., Chen, M., Guo, P., Dai, Z., et al., 2021. clusterProfiler 4.0: A universal enrichment tool for interpreting omics data, Innovation (Camb). 2:100141.

[15] Chamberlain, J. R., Lee, Y., Lane, W. S., Engelke, D. R., 1998. Purification and characterization of the nuclear RNase P holoenzyme complex reveals extensive subunit overlap with RNase MRP, Genes Dev. 12:1678–90.

[16] Lin, J., Ren, J., Gao, D. S., Dai, Y., Yu, L., 2021. The Emerging Application of Itaconate: Promising Molecular Targets and Therapeutic Opportunities, Front Chem. 9:669308.

[17] Mills, E. L., Ryan, D. G., Prag, H. A., Dikovskaya, D., Menon, D., Zaslona, Z., et al., 2018. Itaconate is an anti-inflammatory metabolite that activates Nrf2 via alkylation of KEAP1, Nature. 556:113–117.

[18] Amici, A., Emanuelli, M., Magni, G., Raffaelli, N., Ruggieri, S., 1997. Pyrimidine nucleotidases from human erythrocyte possess phosphotransferase activities specific for pyrimidine nucleotides, FEBS Lett. 419:263–7.

[19] Watanabe, S., Uchida, T., 1995. Cloning and expression of human uridine phosphorylase, Biochem Biophys Res Commun. 216:265–72.

