## Supplemental Data for "Combined single-sample metabolomics and RNAseq reveals a hepatic pyrimidine metabolic response to acute viral infection"

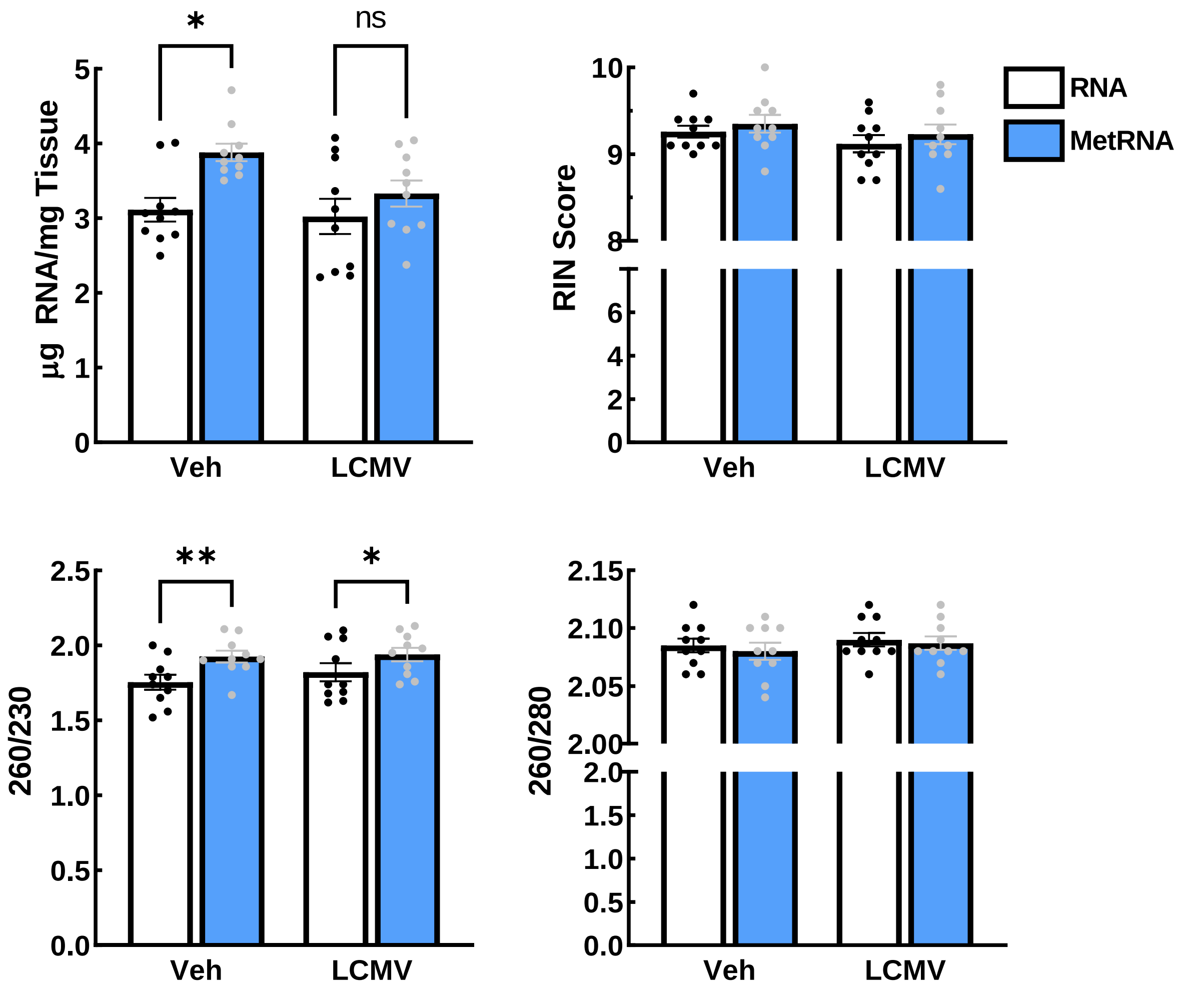


Figure S1. Characterization of the quality and yield of the mRNA after extraction.

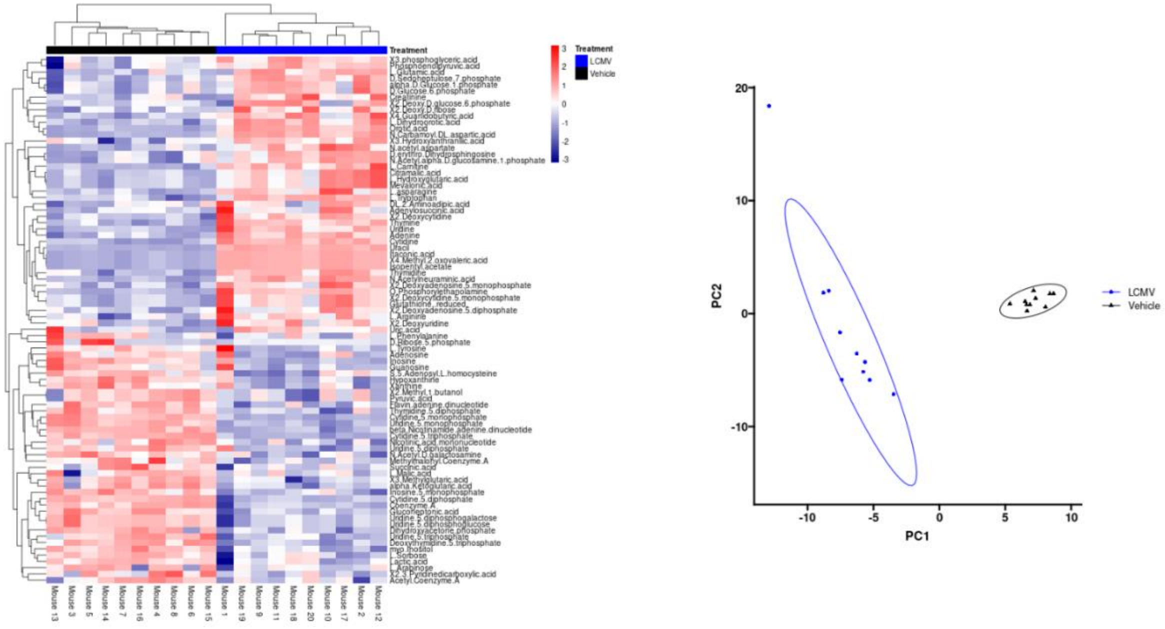


Figure S2. Hepatic metabolomic analysis. Differentially abundant metabolites show strong clustering by group in both a heatmap and principal component analysis.

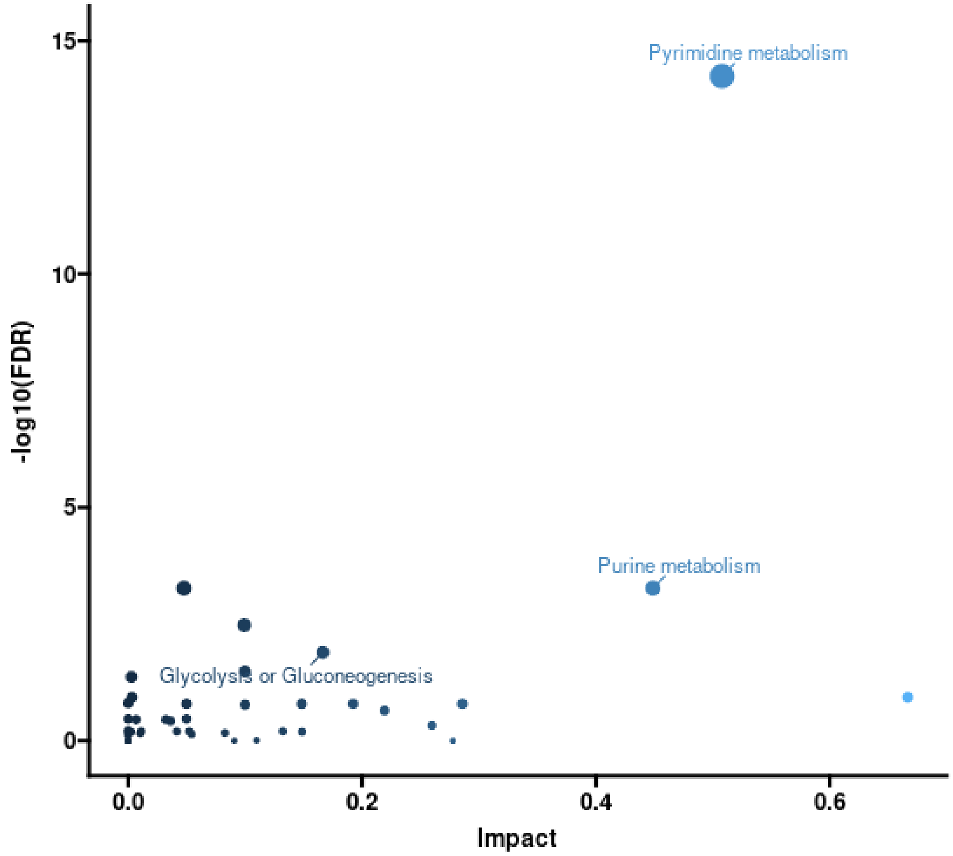


| | Total | Expected | Hits | FDR | Impact |
|---|---|---|---|---|---|
| Pyrimidine metabolism | 39 | 1.8 | 19 | <0.0001 | 0.51 |
| Purine metabolism | 66 | 3.0 | 12 | 0.0005 | 0.45 |
| Glycolysis or Gluconeogenesis | 26 | 1.2 | 6 | 0.013 | 0.17 |
| Pyruvate metabolism | 22 | 1.0 | 5 | 0.033 | 0.1 |
| Citrate cycle (TCA cycle) | 20 | 0.90 | 6 | 0.003 | 0.1 |

Figure S3. Metabolite only pathway analysis.

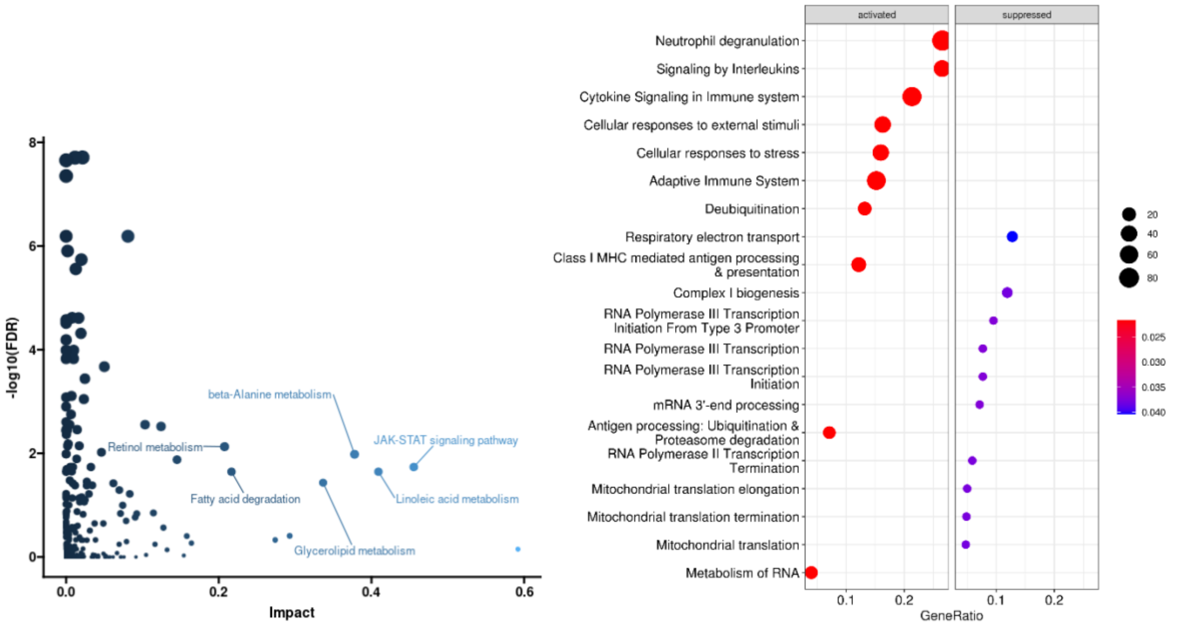


| | Total | Expected | Hits | FDR | Impact |
|---|---|---|---|---|---|
| JAK-STAT signaling pathway | 165 | 14.6 | 26 | 0.02 | 0.46 |
| Linoleic acid metabolism | 50 | 4.4 | 11 | 0.02 | 0.41 |
| beta-Alanine metabolism | 32 | 2.8 | 9 | 0.01 | 0.38 |
| Glycerolipid metabolism | 61 | 5.4 | 12 | 0.04 | 0.34 |
| Fatty acid degradation | 50 | 4.4 | 11 | 0.02 | 0.22 |

Figure S4. Gene only pathway analysis and pathway enrichment from clusterProfiler.

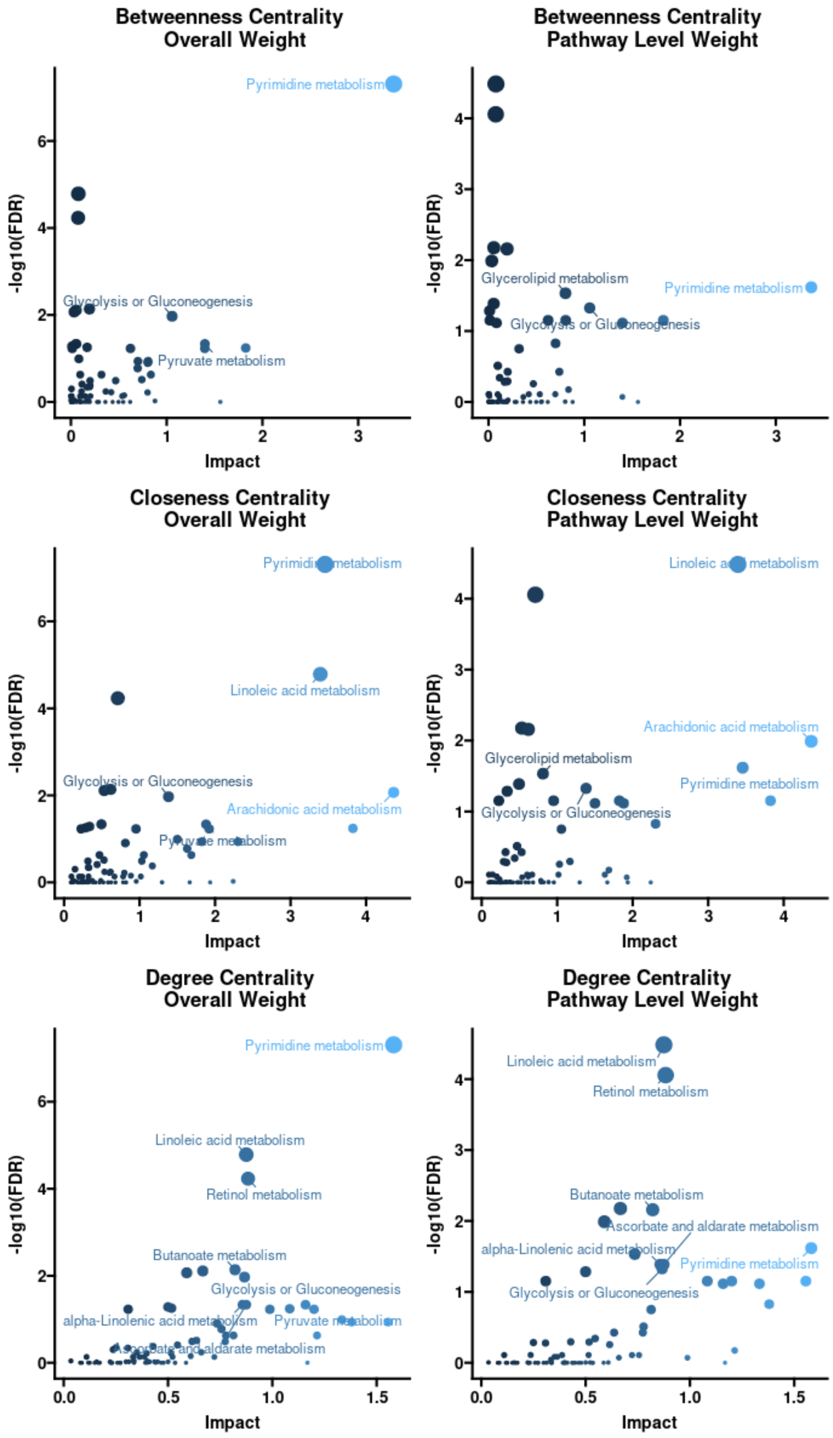


Figure S5. Joint metabolic and transcriptomic enrichment analysis under other degree centralities and weighting methods.
